# Transposable temperate phages promote the evolution of divergent social strategies in *Pseudomonas aeruginosa* populations

**DOI:** 10.1101/593798

**Authors:** Siobhán O’Brien, Rolf Kümmerli, Steve Paterson, Craig Winstanley, Michael A. Brockhurst

## Abstract

Transposable temperate phages randomly insert into bacterial genomes, providing increased supply and altered spectra of mutations available to selection, thus opening alternative evolutionary trajectories. Transposable phages accelerate bacterial adaptation to new environments, but their effect on adaptation to the social environment is unclear. Here we show, using experimental evolution of *Pseudomonas aeruginosa* in iron-limited and iron-rich environments causing differential expression of siderophore cooperation, that transposable phages promoted divergence into extreme siderophore production phenotypes in iron-limited populations. Iron-limited populations with transposable phages evolved siderophore over-producing clones alongside siderophore non-producing cheats. Low siderophore production was associated with parallel mutations in *pvd* genes, encoding pyoverdine biosynthesis, and *pqs* genes, encoding quinolone signaling, while high siderophore production was associated with parallel mutations in phenazine-associated gene clusters. Notably, some of these parallel mutations were caused by phage insertional inactivation. These data suggest that transposable phages, which are widespread in microbial communities, can mediate the evolutionary divergence of social strategies.

## Introduction

Bacteriophages (phages) are viruses that infect and replicate within bacterial cells, and outnumber eukaryotic viruses both in abundance and diversity (Reyes et al 2012). Phages have been found in almost all environments studied so far, including human gastrointestinal (Breitbart et al 2003; Kim et al 2011) and respiratory tracts (Willner et al 2009). While lytic phages rely solely on horizontal transmission to replicate, temperate phages can either transmit horizontally via cell lysis or vertically as prophages integrated into the bacterial chromosome (Little 2005; Ghosh et al 2009). Acquisition of prophages can promote bacterial evolutionary innovation by providing bacteria with new accessory gene functions encoded on the phage genome or via co-option of phage genetic material to form new bacterial traits (e.g. phage tail-derived bacteriocins (Strauch et al 2001)). Alternatively, temperate phages can increase supply and alter the spectra of mutations available to selection through insertional inactivation of bacterial genes by prophage (Harrison & Brockhurst 2017). In particular, transposable phages, which integrate randomly throughout the bacterial genome, can increase genetic diversity and accelerate adaptation of bacterial populations (Davies et al 2016). For instance, the transposable phage LESϕ4 mediated adaptation of *Pseudomonas aeruginosa* to a new environment (artificial sputum medium mimicking the physiochemical properties of sputum in the cystic fibrosis lung) by increasing the supply of positively selected mutations in motility- and quorum sensing-associated genes (Davies et al 2016).

Transposable phages could also affect the evolution of bacterial cooperative interactions, such as siderophore production, by increasing the supply of cheater mutants: genotypes benefiting from but not paying the cost of cooperation (Griffin et al 2004). Siderophores are secreted by many bacterial species in response to iron-starvation (Holden & Bachman 2015). Siderophore production can be cooperative, since the production of siderophores carries a cost to the producing individual (Griffin et al 2004), but the siderophore-iron complex can be taken up by any nearby cell with the appropriate cell-surface receptor. This can make siderophore producers vulnerable to invasion by *de novo* non-producing cheats, who lose the ability to produce siderophores but still retain the ability to use siderophores produced by others (Harrison & Buckling 2005, 2007; O’Brien et al 2013). Thus, similarly to the effect of an increased genomic mutation rate due to inefficient mismatch repair, we would predict that transposable phages would drive the breakdown of siderophore cooperation by increasing the rate and probability of efficient cheats evolving (Harrison & Buckling 2005, 2007).

Here, we examine the impact of the transposable temperate phage LESϕ4 on the evolutionary dynamics of siderophore cooperation in *P. aeruginosa*. Populations were experimentally evolved in either iron-limited or iron-rich culture conditions where cooperative siderophore production is, respectively, required or not required for growth. We observed that, contrary to our prediction, transposable phage did not drive greater breakdown of mean siderophore cooperation at the population level, but did promote greater divergence of siderophore production among clones and coexistence of extreme social strategies within populations under iron-limitation.

## Materials and methods

### a) Strains and culturing conditions

We used *P. aeruginosa* PAO1 as our siderophore-producing wildtype ancestor, and the temperate transposable phage LESϕ4. LESϕ4 inserts randomly into the host genome (Davies et al 2016), and displays high rates of lytic activity in chronic CF lung infections, including being induced into the lytic cycle by clinically relevant antibiotics (James et al 2015).

We confirmed that our PAO1 strain was susceptible to LESϕ4 using a plaque assay: susceptibility was confirmed by a clear plaque on a bacterial lawn formed by phage-mediated lysis. Evolution experiments were performed in casamino acids medium (CAA; 5g Casamino acids, 1.18g K2HPO43H20, 0.25g MgSO_4_7H_2_0 per litre). Media was made iron-limited through the addition of sodium bicarbonate to a final concentration of 20mM, and 100ug ml^-1^of human apotransferrin. Iron-replete media was established by the addition of 20µM FeSO_4‥_We confirmed that our iron rich conditions negated the fitness cost experienced by non-producers under iron-limitation, by growing wildtype and a siderophore non-producing mutant PAO1*ΔPvdDΔPchEF* (harbouring loss of function mutations in the two major siderophores pyoverdine and pyochlin; Ghysels et al 2004) in iron-rich and limited media in a 96 well plate, and measuring optical density (OD) after 24h (iron-rich: non-producers reach higher densities than wildtype, t-test: t_10_=3.7476, p=0.004; iron-limited: wildtype reach higher densities than non-producers: Kolmogorov-Smirnov test: D=1, p=0.005). Wildtype *per capita* pyoverdine production (see (c) below) was also repressed under iron-rich compared with iron-limited conditions (Welch t-test: t_5.13_=236.84, p<0.0001). The fitness advantage experienced by non-producers in iron-rich environments has been attributed to the cost of harbouring the siderophore producing machinery (Griffin et al 2004).

### b) Evolution experiment

We followed real-time evolutionary changes in production of the most costly and efficient siderophore, pyoverdine, over time (Visca et al 2007; Youard et al 2011; Dumas et al 2013), in response to the present and absence of LESϕ4, in iron-rich and iron-limited CAA. To ensure that each population was colonized by a single colony (i.e. relatedness =1 at the first transfer), PAO1 was cultured in 6ml King’s Medium B (KB; (10 g glycerol, 20 g protease peptone no. 3, 1.5 g K_2_HPO_4._3H_2_O, 1.5 g MgSO_4._7H_2_O, per litre) for 24h at 37°C, after which it was diluted with M9 minimal salt solution and grown for 24h on KB agar at 37°C. A single colony was then selected and grown in KB medium at 37°C for 24h, and subsequently used to establish populations.

To initiate the experiment, twelve 30ml universal glass tubes were filled with 6ml iron-limited CAA, and twelve with iron-rich CAA. PAO1 was inoculated to a density of 5 × 10^6^CFU ml^-1^. 10^6^ml^-1^plaque forming units (PFU’s) of LESϕ4 phage was added to six iron-limited and six iron-rich populations (MOI=1), in a full factorial design. Cultures were grown shaken at 180rpm with slightly loose caps, at 37°C.

Every 24h, 1% of each population was transferred to fresh tubes (iron-rich or limited as appropriate). The experiment was conducted for 30 transfers, and every 2nd transfer populations were mixed with 20% glycerol and frozen at −80°C for further analysis. At transfers 10, 15, 20, 25 and 30, populations were 1) plated on KB agar to assess density (colony forming units (CFU) /ml), and 2) tested for the presence of free phage. Free phage was detected in all populations treated with phage throughout the experiment, however phages had no effect on bacterial densities (LMER; *X*^*2*^_*1,6*_= 0.1698, p=0.68, figure S4).

### c) Pyoverdine Assays

Every 10 transfers, populations were diluted and plated on KB agar. Thirty colonies from each population were selected at random, and pyoverdine production quantified for each colony (2160 colonies total). Using sterile toothpicks, individual colonies were transferred to 120µl KB medium in 96 well plates. Plates were incubated static for 24h at 37°C to allow cultures to reach stationary phase. 1% of each culture was then transferred to 180ul iron-limited CAA (siderophore-stimulating conditions), and incubated static for 24h at 37°C. Colonies were analysed for pyoverdine production (RFU) using a pyoverdine-specific emission assay (Ankenbauer et al 1985; Cox & Adams 1985). Briefly, fluorescence of each culture was measured at 460nm following excitation at 400nm, using a Tecan infinite M200 pro spectrophotometer. Optical density (OD) was also measured at 600nm, and the ratio RFU/OD was employed as a quantitative measure of per capita pyoverdine production (Kümmerli et al 2009 JEB).

### d) Sequencing evolved clones

To identify the underlying genetic drivers of siderophore cheating in the iron-limited phage treatment, we specifically selected the two highest pyoverdine producers and two lowest producers from each population evolving in this treatment. Since the proportion of siderophore producers relative to non-producers did not always permit this at the final transfer, clones were isolated from the iron-limited phage treatment at either the 10^th^(Populations 2,5 and 6) or 30^th^(Populations 1,3 and 4) transfer. To test whether observed parallel mutations in these clones are treatment-specific, we next selected 4 clones at random from every population in the remaining treatments at the final timepoint (72 clones from 18 populations).

The Wizard® Genomic DNA Purification kit (Promega) was used to isolated genomic DNA from overnight cultures, according to the manufacturer’s instructions. The quality and quantity of the isolated gDNA was assessed using Nanodrop (Thermo Scientific) and Qubit, respectively. Illumina Nextera XT genomic libraries were prepared by the Centre for Genomic Research, University of Liverpool and 2 × 250 bp paired-end reads generated on an Illumina MiSeq platform. See the electronic supplementary material, SX for further details on sequence data preparation.

Reads were aligned to the PA01 reference (NC_002516.2) using BWA-MEM, duplicates removed and variants detected and filtered using the Genome Analysis Toolkit (GATK) HaplotypeCaller and VariantFiltration tools (McKenna et al 2010) based on GATK filtering recommendations. Mutations common to all clones were removed from dataset, as these likely represented divergence between our PAO1 stock and the reference genome. Only variants passing filtering were included in the final dataset (196 mutations across all clones). A further 75 were mutations associated with pf1 bacteriophage, and subsequently were removed from analysis (see Davies et al 2016). Of the remaining 121 mutations, 12 were synonymous and were excluded from analysis. The remaining 109 mutations comprised our final dataset, consisting of frameshifts (33%), missense mutations (49%), gene deletions (13%) and stop-gained (6%). Evolved clones acquired between 15 and 32 non-synonymous mutations each, with a mean of 16.47 mutations per clone. We further analysed the data to pinpoint the genes in which phage had inserted (SNP calling does not detect phage-mediated mutations; Davies et al 2016). All reads were mapped to the PA01 and phi4 genomes and reads retained where one member of a read pair mapped to PA01 and the other to phi4. From these, reads mapping to the PA01 genome were counted within 1kbp moving windows and potential insertion sites further inspected in IGV manually (Thorvaldsdóttir et al 2013). Phage insertion sites were identified in all 48 phage-evolved clones isolated from 12 populations.

### e) Statistical analysis

All data were analysed using R version 2.15.1 (R Core Team, 2016). We used a t-test (for normally distributed data and equal variances) and Kolmogorov Smirnov test (non-normal distribution and unequal variances) test to compare Malthusian growth rates (*m*) of each strain under iron rich and iron limited conditions, respectively. Malthusian growth rate (*m*) was quantified as ln(Final density/Starting density) (Lenski et al 1991). To investigate any effect of iron-limitation and/or phage in shaping pyoverdine production, we used linear mixed effects models, assigning per capita pyoverdine as the response variable, iron, phage and transfer number (including 2- and 3-way interactions) as fixed explanatory variables, and population ID and transfer number as random explanatory variables as follows:

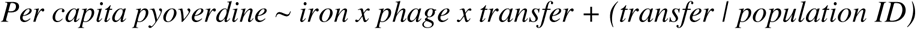

Variance (σ^2^) in pyoverdine production at transfer 10, 20 and 30 was calculated for 30 colonies per population. To investigate any effect of iron-limitation and/or phage in shaping within population variance in pyoverdine production, we used linear mixed effects models, assigning log (population variance) as the response variable, iron, phage and transfer number (including 2- and 3-way interactions) as fixed explanatory variables, and population ID and transfer number as random explanatory variables as follows:

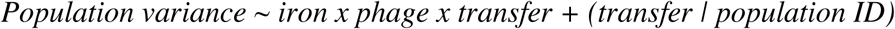

Nonproducers were classified as those colonies producing <5% ancestral pyoverdine levels (<6069.25 RFU). Overproducing colonies were in the 95^th^ percentile (> 146,590 RFU) for per capita pyoverdine production. Ancestral per capita pyoverdine was 121,385 RFU. To determine whether the numbers (counts) of overproducing colonies per population was influenced by treatment, we performed a generalized mixed effects model, with a poisson error structure as follows:

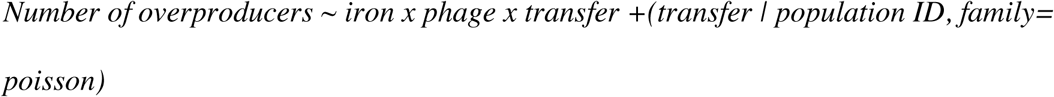

The effect of phage and iron on population density (CFU/ml) was analyzed using linear mixed effects models, assigning population CFU/ml as the response variable, iron and phage (including 2-way interaction) as fixed explanatory variables, and population ID and transfer number as random explanatory variables as follows:

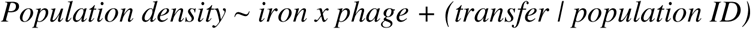

Finally, after ensuring high and low producers did not differ in the variance of nonsynonymous mutations (permutational ANOVA, permutation test:

*F*_1,10_= 0.5373, *P* =0.491), we performed a Permutational Multivariate Analysis of Variance using the adonis function in R to assess mutational differences between phenotypes.

## Results

### a) The evolutionary dynamics of pyoverdine production

To obtain a quantitative measure of siderophore production, we calculated *per capita* pyoverdine production for thirty colonies per population every 10th transfer for 30 transfers. Iron limitation reduced *per capita* pyoverdine production, but this effect weakened over time (LMER; iron x transfer interaction, *X*^*2*^_*1,9*_= 8.1656, p=0.004, figure 1, figure S3). To a lesser extent, transposable phage presence also reduced *per capita* pyoverdine production, but again, this effect became weaker over time (LMER; phage x transfer interaction, *X*^*2*^_*1,9*_= 4.2632, p=0.03, figure 1, figure S3). The degree to which iron limitation affected pyoverdine production was not influenced by the presence of phage over the course of the experiment, or *vice versa* (LMER; non-significant iron x phage x transfer interaction, *X*^*2*^_*1,11*_= 0.0003, p=0.9, figure 1, figure S3).

**Figure 1:**
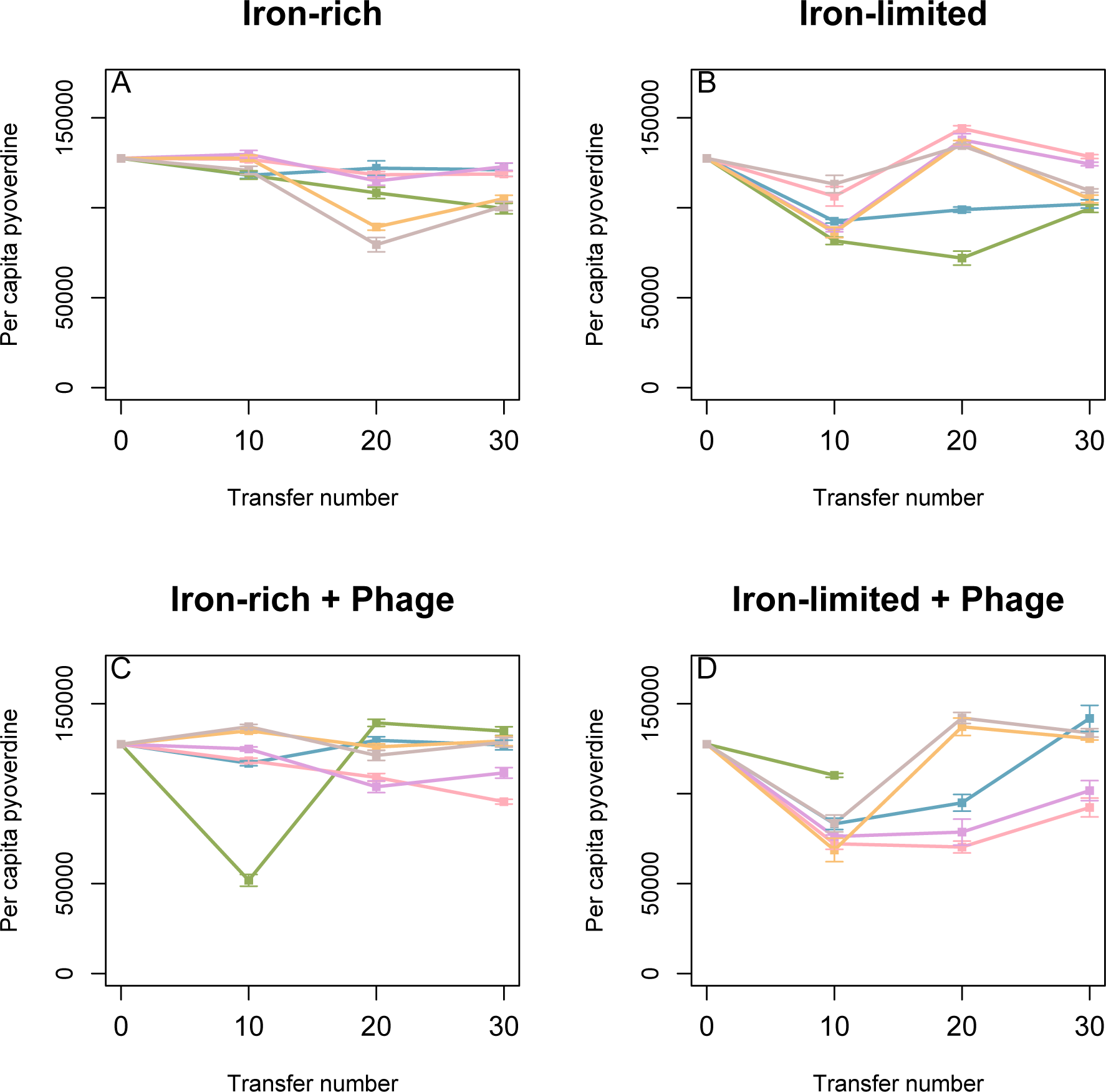
Changes in per capita pyoverdine production over the course of 30 transfers for 24 populations assigned to one of the following treatments A) iron-rich, B) iron-limited C) iron-rich & phage, D) iron-limited & phage. Iron limitation reduced per capita pyoverdine, but this effect was less strong over time (LMER; iron x transfer interaction, *X*^2^_*1,9*_= 8.1656, p=0.004). To a lesser extent, phage presence also reduced *per capita* pyoverdine production, but again, this effect was reduced over the course of the experiment (LMER; phage x transfer interaction, *X*^2^_*1,9*_= 4.2632, p=0.03). The extent to which iron limitation influences pyoverdine production was not influenced by the presence of phage over the course of the experiment, and *vice versa* (LMER; non-significant iron x phage x transfer interaction, *X*^2^_*1,11*_= 0.0003, p=0.9). Data show means 30 isolated colonies per population +/-SEM’s.

Although transposable phages did not affect mean pyoverdine production under iron limitation, populations with temperate phages did show greater variation between clones in their pyoverdine production in these populations. Within-population variance in pyoverdine production was highest in iron-limited populations evolving with phages (LMER; phage x iron interaction, *X*^*2*^_*1,9*_=12.22, p=0.0004, figure 2, figures S1-S3), irrespective of time (LMER non-significant phage x iron x transfer interaction *X*^*2*^_*1,11*_=0.4591, p=0.4981, figure 2, figures S1-S3). Neither iron-limitation nor phage presence alone influenced within-population variance (LMER; iron effect *X*^*2*^_*1,6*_= 1.19, p=0.27; phage effect *X*^*2*^_*1,6*_= 2.99, p=0.08, figure 2, figures S1-S3).

**Figure 2:**
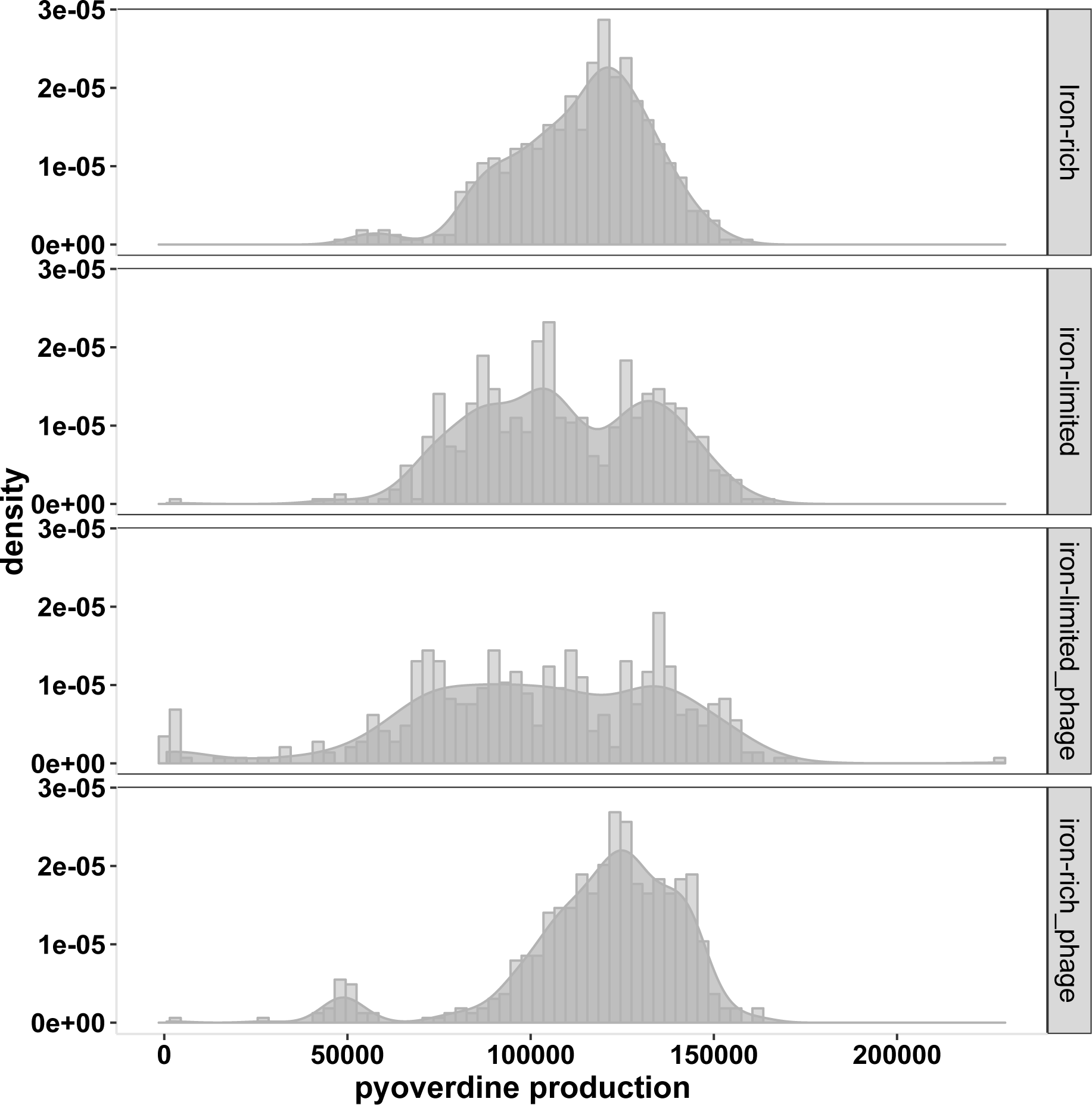
Density histogram illustrating variation in pyoverdine production within each treatment. Data are per capita pyoverdine production for 480-540 isolated colonies per treatment (pooling all replicates and timepoints within a treatment). Within-population variance in pyoverdine production increased only under iron-limitation and in the presence of phage (LMER; phage x iron interaction, *X*^2^_*1,9*_=12.22, p=0.0004), irrespective of time (LMER non-significant phage x iron x transfer interaction *X*^2^_*1,11*_=0.4591, p=0.4981). Neither iron-limitation nor phage presence alone influenced within-population variance (LMER; iron effect *X*^2^_*1,6*_= 1.19, p=0.27; phage effect *X*^2^_*1,6*_= 2.99, p=0.08).

To determine whether this higher variance was driven by increased numbers of pyoverdine over-producers, non-producers, or coexistence of both, we analysed the number of non- and overproducing isolates in each treatment. Numbers of overproducing isolates increased in the presence of phage over time (phage x timepoint interaction: *X*^2^_1,7_= 5.2866, p=0.022), and this was not significantly influenced by iron availability (effect of iron: *X*^2^_1,7_= 5.2866, p=0.4). Nonproducing isolates (n=17) were identified throughout the experiment, predominantly (in 15/17 cases) in iron-limited populations evolving with phages (Table S1). By the final transfer, nonproducing mutants had been observed at least once in 5/6 populations from the iron-limited treatment with phages. Together, this suggests that the increased variation in pyoverdine production observed in the iron limited populations evolved with phages was driven by increased numbers of pyoverdine nonproducers in this treatment in addition to increased numbers of overproducers driven by the presence of phage *per se.*

### b) Genome sequencing of evolved clones

To examine the genetic bases of the observed divergence in pyoverdine production phenotypes observed in iron-limited populations evolved with phages, we obtained whole genome sequences for the two highest- and the two lowest-pyoverdine-producing clones from each population (i.e. 24 clones; 4 clones each from 6 populations). Comparing the highest- (n=12) and lowest-producing clones (n=12), we sought to identify mutations distinguishing these classes. First, we analysed all nonsynonymous mutations, including SNPs and indels together with genes affected by phage insertional inactivation mutations. High and low producers did not differ in the variance of nonsynonymous mutations (permutational ANOVA, permutation test: *F*_1,10_= 0.5373, *P* =0.491), but the loci affected by non-synonymous mutations did differ between high and low producers (permutational ANOVA, permutation test: *F*_1,10_= 2.1429, *P* =0.014, Figure 3, Table S2).

**Figure 3:**
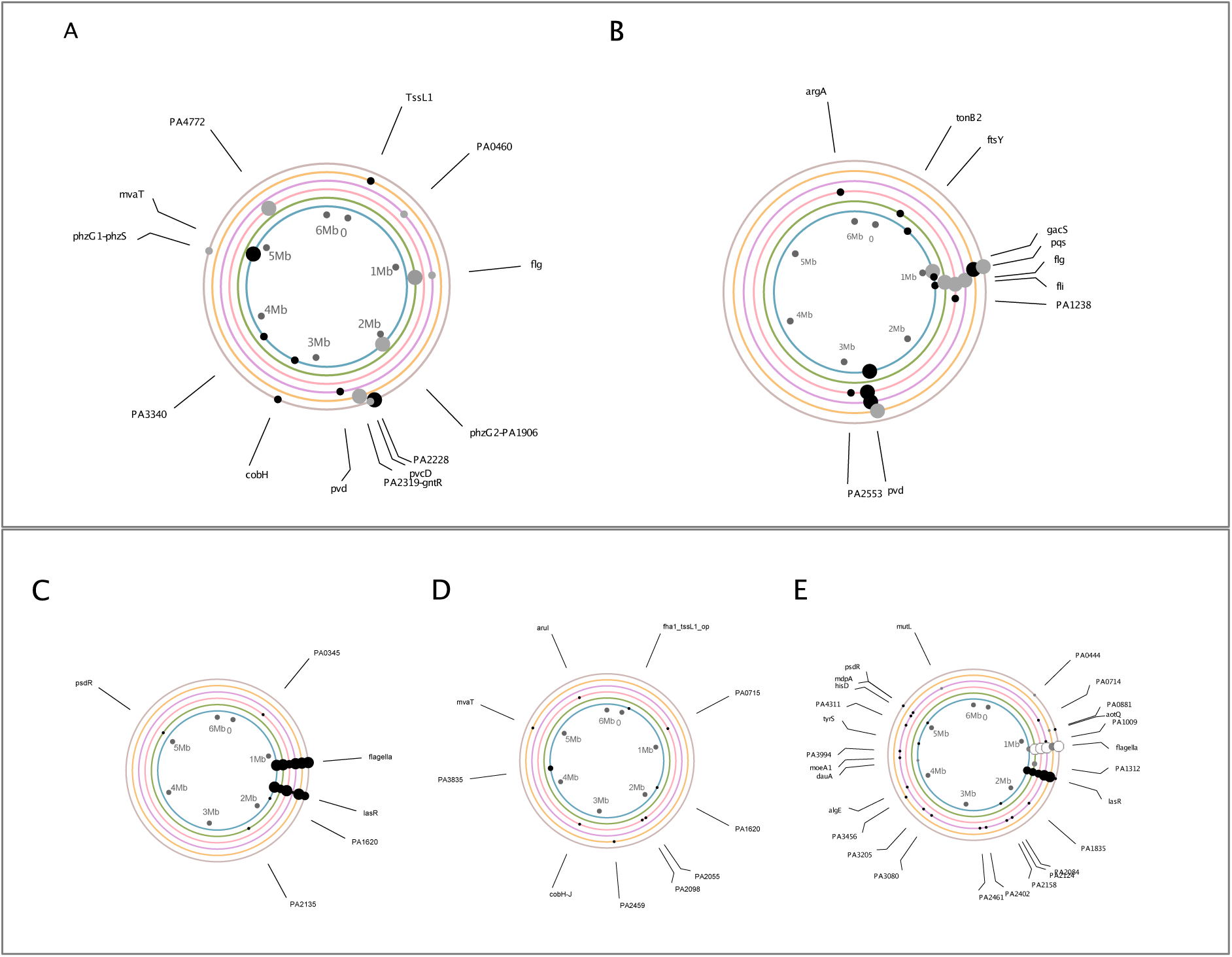
Genetic loci under positive selection in high (**A**) and low (**B**) pyoverdine producing clones evolving with phages in iron-limited media. Each concentric circle represents a replicate population in either high (**A**) or low (**B**) producers. Concentric circles correspond to populations 1-6, from the innermost to outermost circle, respectively. Positions around each concentric circle correspond to positions around the PAO1 published and annotated chromosome. Small black dots around these circles indicate the occurrence of an indel or SNP, grey dots represent phage integration events in those regions, and white dots indicate both. C, D, and E correspond to control populations evolving under iron-rich (C), iron-limited (D) and iron-rich with phage (E) conditions. Four colonies were selected in total per population: Dot size corresponds to the number of colonies in which a given mutation was observed. When two genes are mentioned, the mutation is intergenic. A complete list of mutations can be found in Tables S2 and S3.

To identify loci likely to have been under divergent positive selection between high and low producers, we looked for evidence of phenotype-specific parallel evolution, i.e. pathways targeted by mutation in multiple clones isolated from independently evolving replicate populations that occurred exclusively in either high or low pyoverdine producers. In low producing clones, we observed parallel mutations in quorum-sensing associated loci (*pqsA, pqsR*; 3/6 populations; 5 clones, Figure 3A,B, Figure 4, Table S2) and pyoverdine biosynthesis associated loci (*pvdA, pvdD, pvdI, pvdS* binding site; 4/6 populations; 8 clones, Figure 3A,B, Figure 4, Table S2). Notably, two of these parallel mutations were caused by prophage-mediated insertional inactivation into *pqsA* and *pvdD* genes, respectively. Furthermore, while clones with *pqs* mutations produced less pyoverdine than the ancestor (figure 4), clones harbouring both *pvd* and *pqs* mutations did not produce any detectable pyoverdine (figure 4).

**Figure 4:**
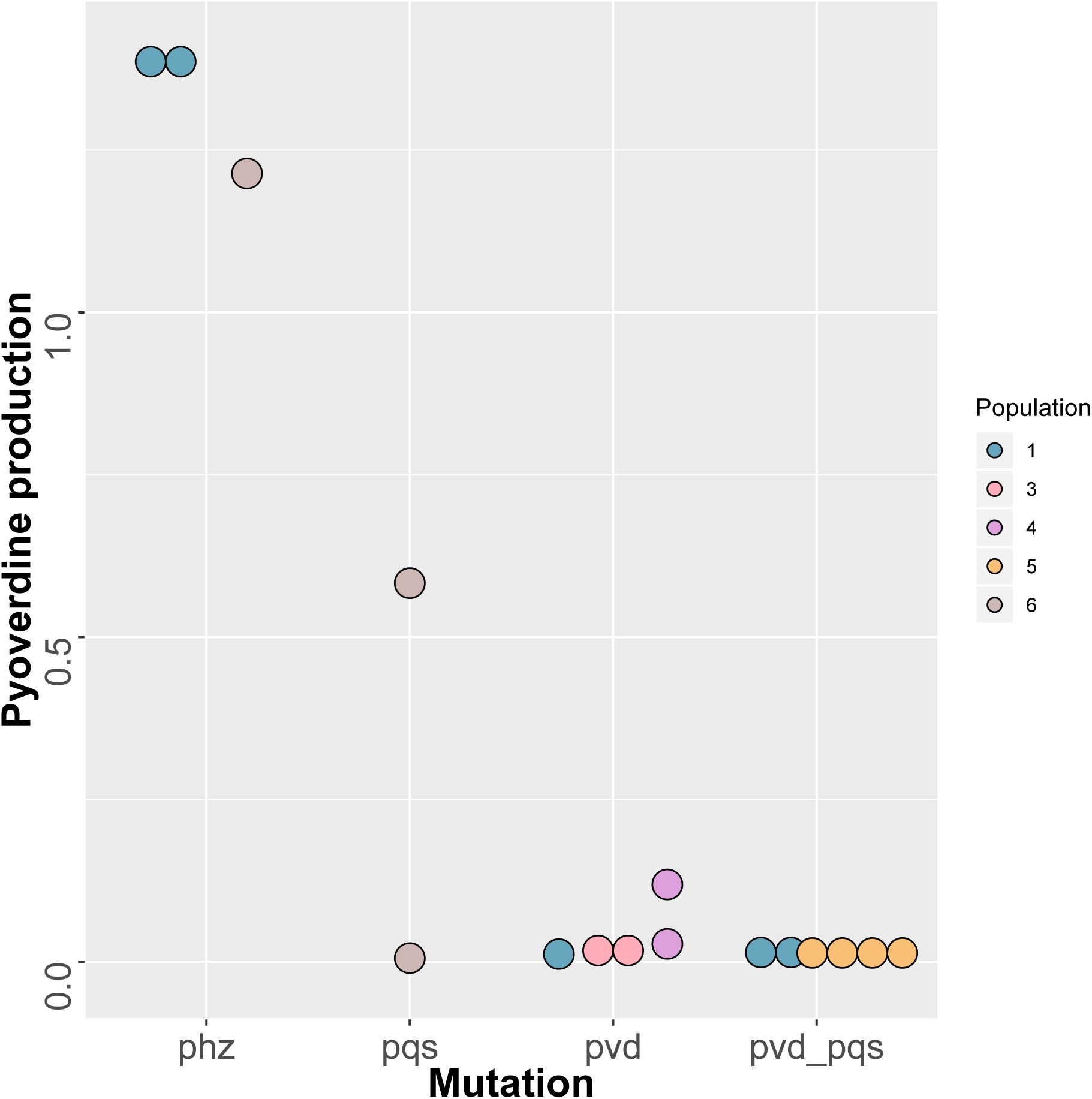
Pyoverdine production relative to ancestor for all clones harbouring mutations in *phz, pvd* or *pqs* associated loci in iron-limited population evolving with phage. Clones are colour coded based on the population from which they originate. *Phz* mutants produce higher pyoverdine relative to the ancestor (output > 1), while *pvd* and *pqs* mutations are associated with reduced pyoverdine production (output < 1).

Exclusive to the highest-producing clones, we observed parallel mutation of a phenazine (*phz*)-associated intergenic region in 2/6 populations (3 clones), all of which were caused by prophage insertion (Figure 3A,B, Figure 4, Table S2). These patterns of distinct parallel evolution in high and low producers are suggestive of divergent selection in the iron-limited populations evolving with phages, and, furthermore, shows that prophage-mediated insertional inactivation mutations contributed to the response to this divergent selection. In addition, we observed shared targets of parallel evolution in flagella associated genes (*flgE, flgF, flgG, flgJ, fliF, fliI*) in both highest- (2/6 populations; 3 clones) and lowest-producer clones (4/6 populations; 7 clones). This suggests that loss of flagellar motility was beneficial *per se* in this well-mixed laboratory scenario, and, moreover, since all but one of these mutations were caused by prophage insertional inactivation, that transposable phage mediated this response to selection.

Next, to determine whether the identified targets of parallel evolution were specific to the iron-limited populations evolved with phages, we obtained whole genome sequences for 4 randomly chosen evolved colonies from each of the replicate populations from the other treatments (Table S3). Because, in these treatments, variation among clones in siderophore production was less extreme, colonies from these populations were chosen at random (instead of selecting high and low producers). Importantly, we never observed mutations in *pqs, pvd* or *phz* associated loci in any of these other treatments. This suggests that mutations in *pqs, pvd* or *phz* associated loci predominantly occurred in iron-limited populations evolved with phages, indicating that these populations followed an evolutionary trajectory distinct from both iron-rich populations and iron-limited populations evolved without phages.

Populations evolved in iron-rich environments underwent highly parallel evolution of the quorum sensing master regulator *lasR*, with 12/12 populations acquiring nonsynonymous SNPs or indels irrespective of phage presence or absence. *lasR* mutations were never caused by prophage insertional inactivation and were never observed in the iron-limited environment, suggesting that loss of *lasR* was an adaptation to iron-rich conditions and was unaffected by the presence or absence of phages. Mutation of flagella-associated loci was observed in all iron-rich populations irrespective of whether phages were present or absent, confirming that loss of flagellar motility is likely to be adaptive *per se* in this well-mixed laboratory environment. When phage was present, prophage-mediated insertional inactivation was the primary mode of flagella-associated mutation, occurring in all 6 replicate populations. Interestingly, under iron limitation, the increased mutational supply provided by transposable phage insertion appears to have promoted loss of flagellar motility: Mutation of flagella-associated loci was not observed in iron-limited populations evolved without phage, in contrast to iron-limited populations evolved with phage where such mutations were common and frequently associated with prophage insertional inactivation (described above).

## Discussion

Transposable phages increase the supply of mutations and thereby can accelerate bacterial adaptation to a new environment (Davies et al 2016). Here, we show that as well as enhancing the response to abiotic environmental selection, transposable phages can also mediate evolutionary responses to the social environment. Populations evolving with phages in an iron-limited environment requiring cooperative siderophore production followed a distinct evolutionary trajectory compared both to populations in the same environment evolved without phages and to populations evolved with or without phage in an iron-rich environment not requiring cooperation. Specifically, within iron-limited populations evolved with phage we observed greater variation in pyoverdine production among clones. This was caused by a combination of more overproducing mutants emerging in populations evolving with phage (irrespective of iron availability), and the emergence of more under-producing and non-producing mutants specifically in iron-limited populations evolving with phage. Our results suggest therefore that transposable phage promoted the evolution of divergent social strategies.

We examined the genetic basis of divergence between the highest- and lowest-producing clones that evolved in the iron-limited treatment with phage. In the lowest-producer class, we observed that pyoverdine synthesis (*pvd*) and regulators of PQS quorum sensing *(pqs)* associated loci were repeatably targeted by parallel mutations in independent replicate lines. Mutation of *pvd*- loci encoding pyoverdine synthesis can cause reduced pyoverdine production (Lamont & Martin 2003; Kümmerli et al 2015; Granato et al 2018, this study (figure 4) and are likely to explain low pyoverdine production in some clones. However, *pvd* mutations were not universal among low pyoverdine producers, and one of the evolved clones from the highest-producer class also carried a *pvd* mutation and maintained ancestral levels of pyoverdine production (Table S2). It is likely therefore, that *pqs* mutations also played a role in the evolution of reduced pyoverdine production. Consistent with this, *P. aeruginosa pqsR* null-mutants produce reduced pyoverdine compared with wildtype (Popat et al 2017; this study (figure 4)), and we could not detect any pyoverdine production in clones harbouring both *pqs* and *pvd* mutations (figure 4). Interestingly, the PQS signal molecule acts as an iron-chelator, so loss of PQS quorum sensing acts to increase the relative availability of iron, reducing the need for siderophore production (Diggle et al 2007; Popat et al 2017). Hence, *pqs* mutants are not impaired in making pyoverdine, but respond phenotypically to increased iron in their environment.

Exclusive to the highest-producing clones, we observed parallel mutation of a phenazine (*phz*)-associated intergenic region, mediated by prophage insertion between *phzG2-PA1906*, and *phzG1-phzS*. *P. aeruginosa* operons phzA1-G1 and phzA2-G2 are involved in biosynthesis of the pyocyanin precursor, phenazine-1-carboxylic acid (PCA). The intergenic region *phzG2-PA1906* contains a putative transcriptional terminator 30bp downstream of the *phzG2* stop codon, and the intergenic region directly downstream of the *phz1* cluster (*phzG1-phzS*) contains a ribosome binding site (Mavrodi et al 2001). Hence, it is likely that these mutations impede both stability and efficiency of the translated phenazine product.

Since pyocyanin increases iron-availability by reducing Fe^3+^ to bioavailable Fe^2+^(Cox 1986) it is possible that high pyoverdine production in these mutants is a response to reduced iron availability caused by reduced levels of the pyocyanin precursor PCA (Granato et al 2018). Interestingly, *pqs* mutations in low pyoverdine producers were always found in the same populations as clones with *phz* mutations – hence, it is possible that *pqs* mutations are a response to, or a driver of, *phz* mutations in coexisting clones.

Adaptation to the iron-rich environment was associated with acquisition of nonsynonymous SNPs or indels in *LasR*, the master regulator of acyl-homoserine lactone (AHL) quorum sensing. This pattern was irrespective of phage, occurring both in their presence or absence, and the mutations were not mediated by prophage insertion. Loss of AHL quorum sensing is a common adaptation to the CF lung (Smith et al 2006; Fothergill et al 2007; LaFayette et al 2015; Davies et al 2016), the *Caenorhabditis elegans* gut (Granato et al 2018; Jansen et al 2015) as well as laboratory media (O’Brien et al 2017; Granato et al 2018). Similarly, loss of *lasR* in CAA is unsurprising – *lasR* is costly and not required in CAA (Sandoz et al 2007; Diggle et al 2007; Darch et al 2012; Ghoul et al 2014). Ghoul et al (2014) demonstrated that a *lasR* mutant grows faster than the wildtype producer in CAA, both when grown alone and in direct competition – showing that mutations in this region are beneficial *per se*. However, while we found *lasR* mutations in 12/12 populations evolving under iron-rich conditions, this was never observed under iron-limitation. This may be because *lasR* expression is enhanced by iron-limitation (Bollinger et al 2001), suggesting that functions regulated by LasR are beneficial in this environment. An alternative explanation is that iron limited populations failed to undergo *lasR* mutations because their densities were lower and evolutionary potential decreased compared with iron rich (Figure S4). However, even under iron limited conditions, populations reached 10^8^−10^9^ CFU’s/ml, which is roughly tenfold higher than that observed in CF artificial sputum – conditions under which *lasR* mutations are commoly observed (Davies et al 2016). Hence, density alone is unlikely to explain the lack of *lasR* mutation under iron-limited conditions.

Loss of flagellar motility appears to have been a common response to selection in our shaken liquid cultures, and mutations in flagella-associated genes were frequently caused by prophage-mediated insertional inactivation mutations. Flagella loss was observed in every (12/12) iron-rich population (irrespective of phage) and was mediated by phage insertional inactivation in every case when phage was present. Under iron-limited conditions, flagella loss was less common (4/12 populations; 10 clones) and only observed in the presence of phages, where in 9/10 cases this was mediated by prophage insertional inactivation, suggesting that transposable phage promoted the loss of flagellar motility by increasing mutational supply. Lim et al (2012) found that in a closely related species, *Pseudomonas fluorescens*, flagella expression was reduced under iron-limitation. Hence, it is possible that the selective benefit of losing flagella is reduced in iron-limited compared with iron-rich conditions. However, mutations in flagella associated loci have been previously observed in *P. aeruginosa* evolving in an iron-limited liquid culture environment after 150 generations (Kümmerli et al 2015), and in iron-limited artificial sputum medium (Davies et al 2016). In our study, population densities were lower under iron-starvation so the reduced mutation supply may explain the lack of flagella mutations under iron-limitation, which was alleviated by the increased mutational supply afforded by transposable phage (Figure S4).

Our results also have applied relevance: *P. aeruginosa* is a common cause of chronic lung infections in cystic fibrosis and bronchiectasis where an initially clonal population undergoes rapid and extensive evolutionary diversification. Analysis of the sputum of cystic fibrosis patients infected with *P. aeruginosa* shows that even within a single sample, high diversity exists in many virulence-associated social phenotypes, such as, pyoverdine, LasA protease and pyocyanin secretions with coexistence of overproducers alongside nonproducers, as we observed in iron-limited populations evolved with phages (Mowat et al 2011; O’Brien et al 2017). Lung sputum is typically iron-limited and siderophores are thought to play an important role in growth, although iron becomes more available over time as the lung tissue deteriorates, and siderophore nonproducers gradually increase in frequency (Andersen et al 2015; Tyrrell & Callaghan 2016). Crucially, temperate phages, including the transposable phage LESϕ4 used in this study, are active and undergo lysis in the CF lung (James et al 2015), suggesting that transposable phage may play a role in the generation of genetic diversity in chronic infections.

## Acknowledgments

This work was funded by a fellowship from the Centre for Chronic Diseases and Disorders at the University of York to SOB (2015-2017), a Phillip Leverhulme Prize to MAB, the European Research Council under the Grant Agreement No 681295 (RK) and the Swiss National Science Foundation No. 31003A_182499 (RK). We would like to acknowledge support from staff at the Centre for Genomic Research, University of Liverpool.

