## Supplementary Material for "Transposable temperate phages promote the evolution of divergent social strategies in *Pseudomonas aeruginosa* populations"

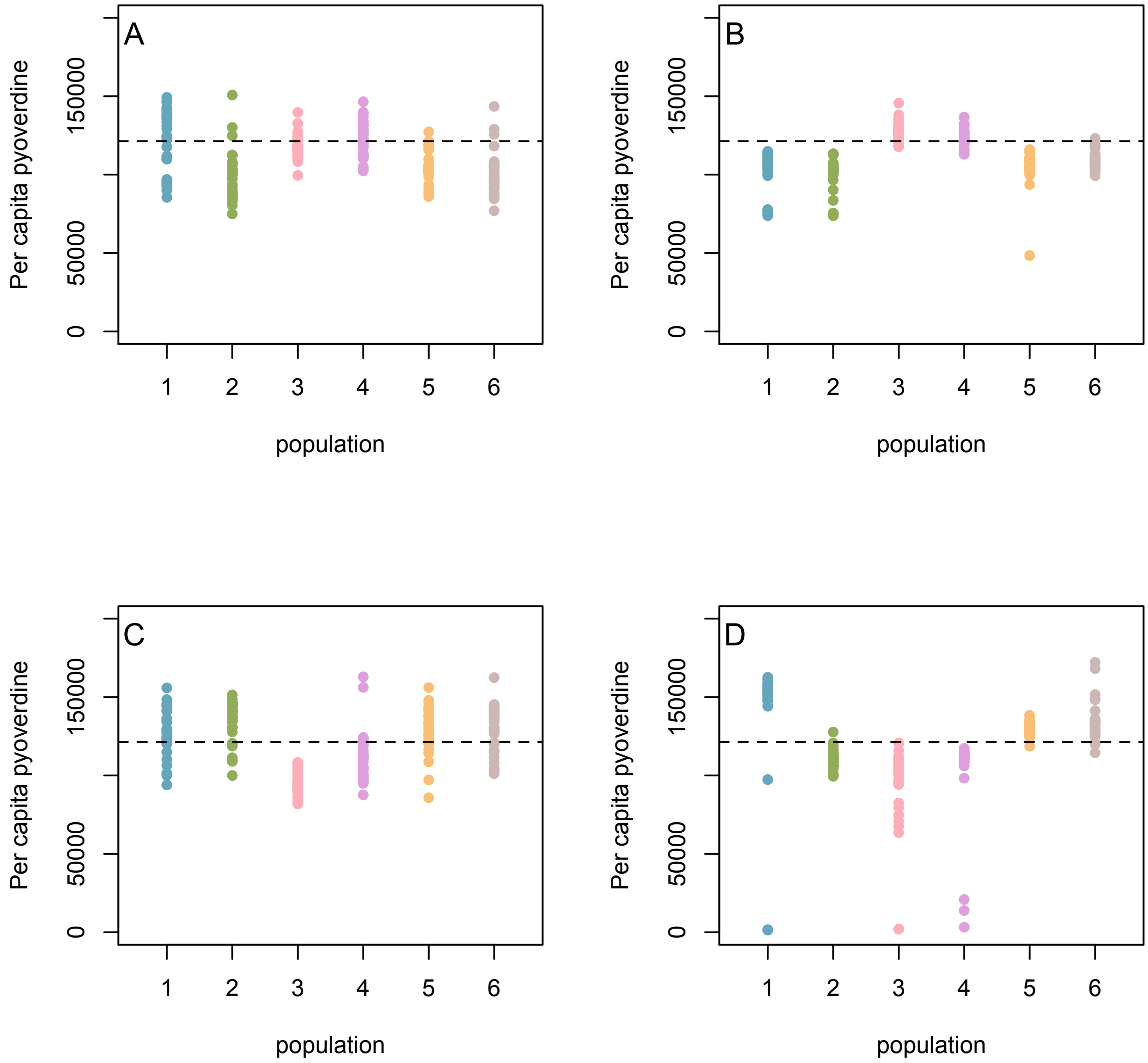

**Figure S1**

Endpoint per capita pyoverdine production for 30 isolated colonies (indicated here as individual dots) per evolving population. Horizontal line indicates ancestral pyoverdine production. Populations were evolved under four treatments: A) iron-rich, B) iron-limited C) iron-rich & phage, D) iron-limited & phage. Within-population variance in pyoverdine production increased only under iron-limitation and in the presence of phage (LMER; phage x iron interaction, $X^2_{1,9}$=12.22, p=0.0004), irrespective of time (LMER non-significant phage x iron x transfer interaction $X^2_{1,11}$=0.4591, p=0.4981).

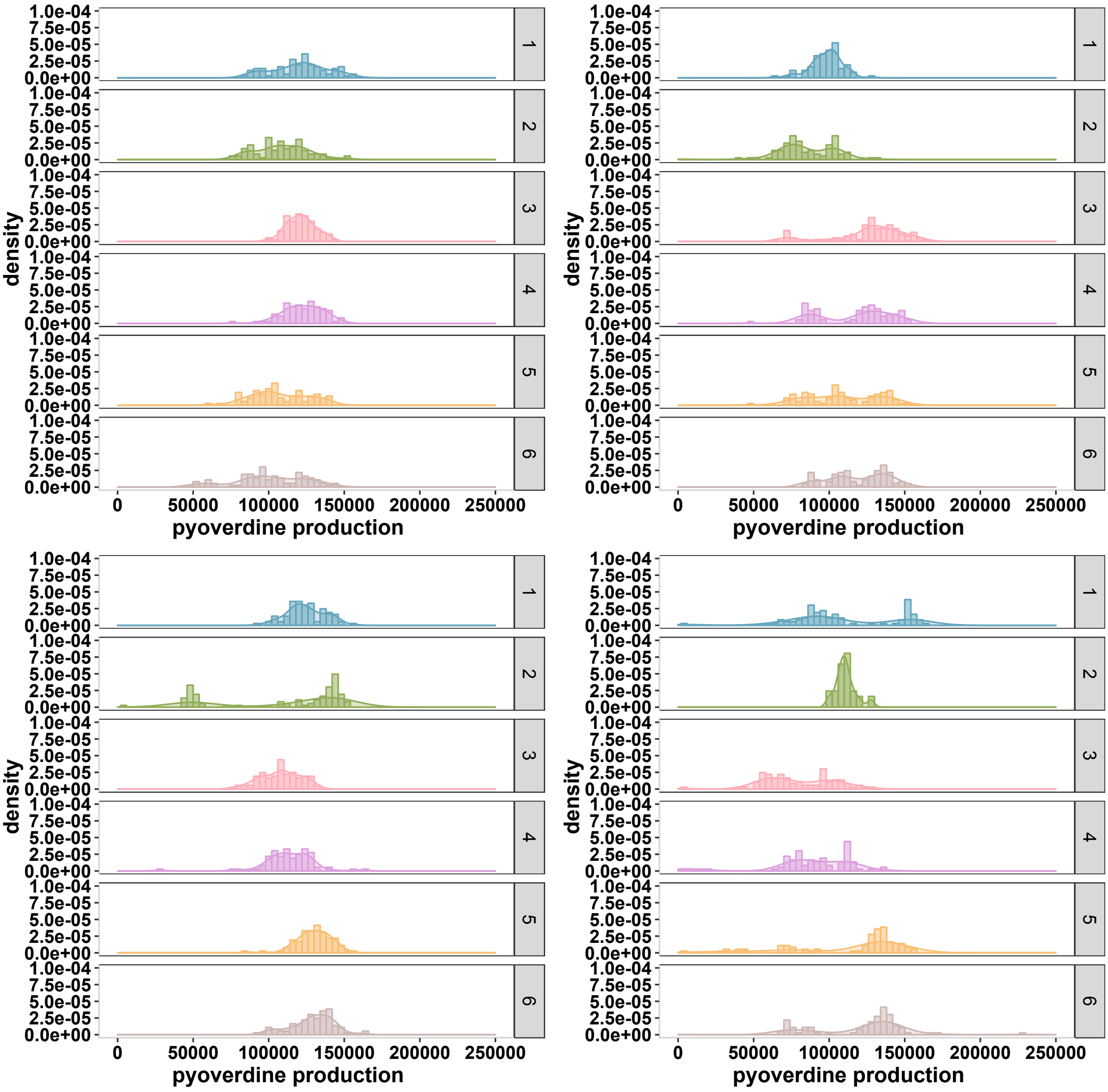

**Figure S2**

Density histogram illustrating variation in pyoverdine production within each evolving population (labeled 1-6 on right side panel). Data are per capita pyoverdine production for 30-90 isolated colonies per treatment (pooling all timepoints for each treatment). Within-population variance in pyoverdine production increased only under iron-limitation and in the presence of phage (LMER; phage x iron interaction, $X^2_{1,9}$=12.22, p=0.0004), irrespective of time (LMER non-significant phage x iron x transfer interaction $X^2_{1,11}$=0.4591, p=0.4981).

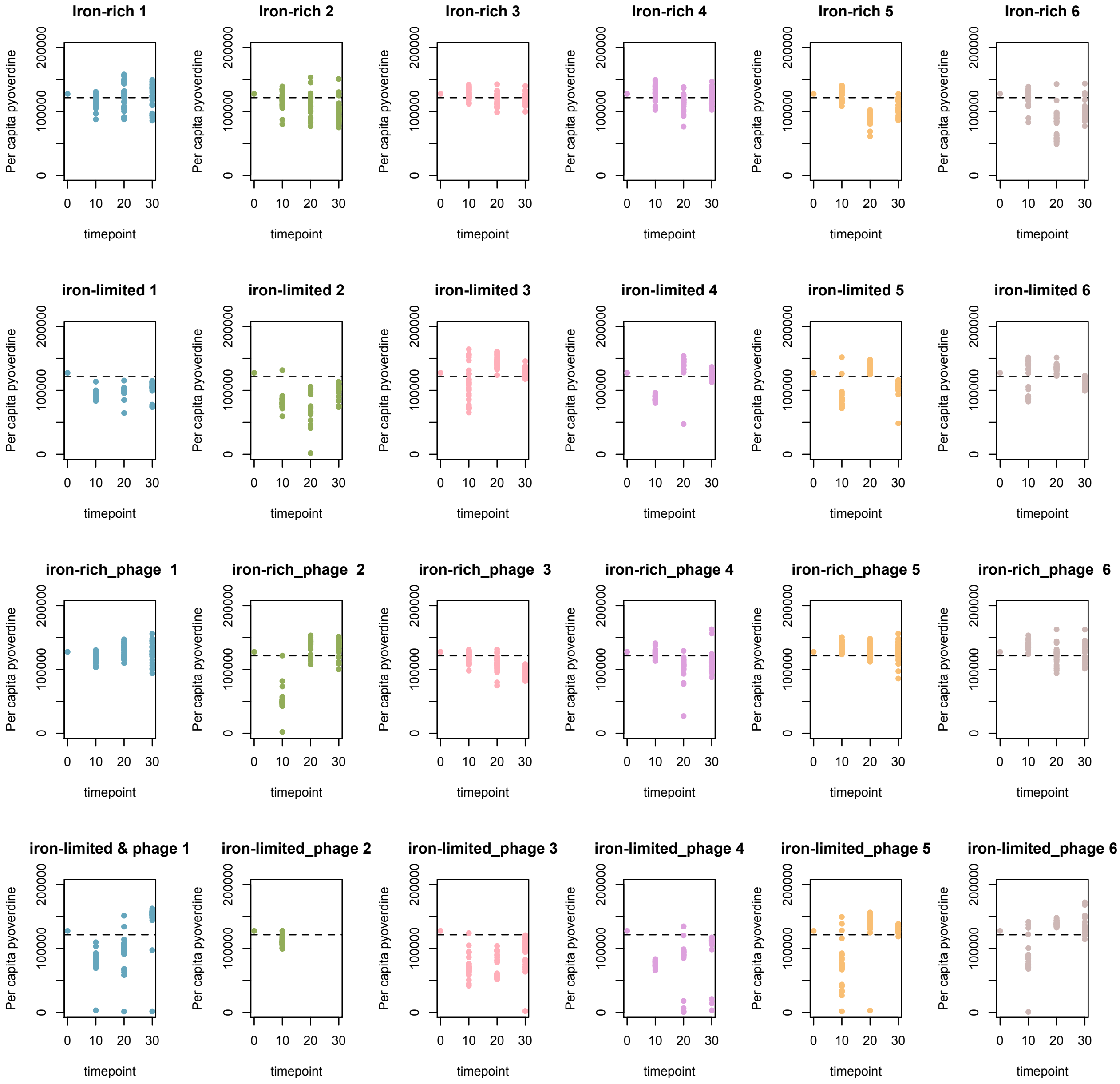

**Figure S3**

Per capita pyoverdine production for every isolated colony through time per evolving population. Horizontal line indicates ancestral pyoverdine production. Iron limitation and phage presence both independently reduced *per capita* pyoverdine, but this effect was less strong over time (LMER; iron x transfer interaction, $X^2_{1,9}$= 8.1656, p=0.004, phage x transfer interaction, $X^2_{1,9}$= 4.2632, p=0.03). Within-population variance in pyoverdine production increased only under iron-limitation and in the presence of phage (LMER; phage x iron interaction, $X^2_{1,9}$=12.22, p=0.0004), irrespective of time (LMER non-significant phage x iron x transfer interaction $X^2_{1,11}$=0.4591, p=0.4981).

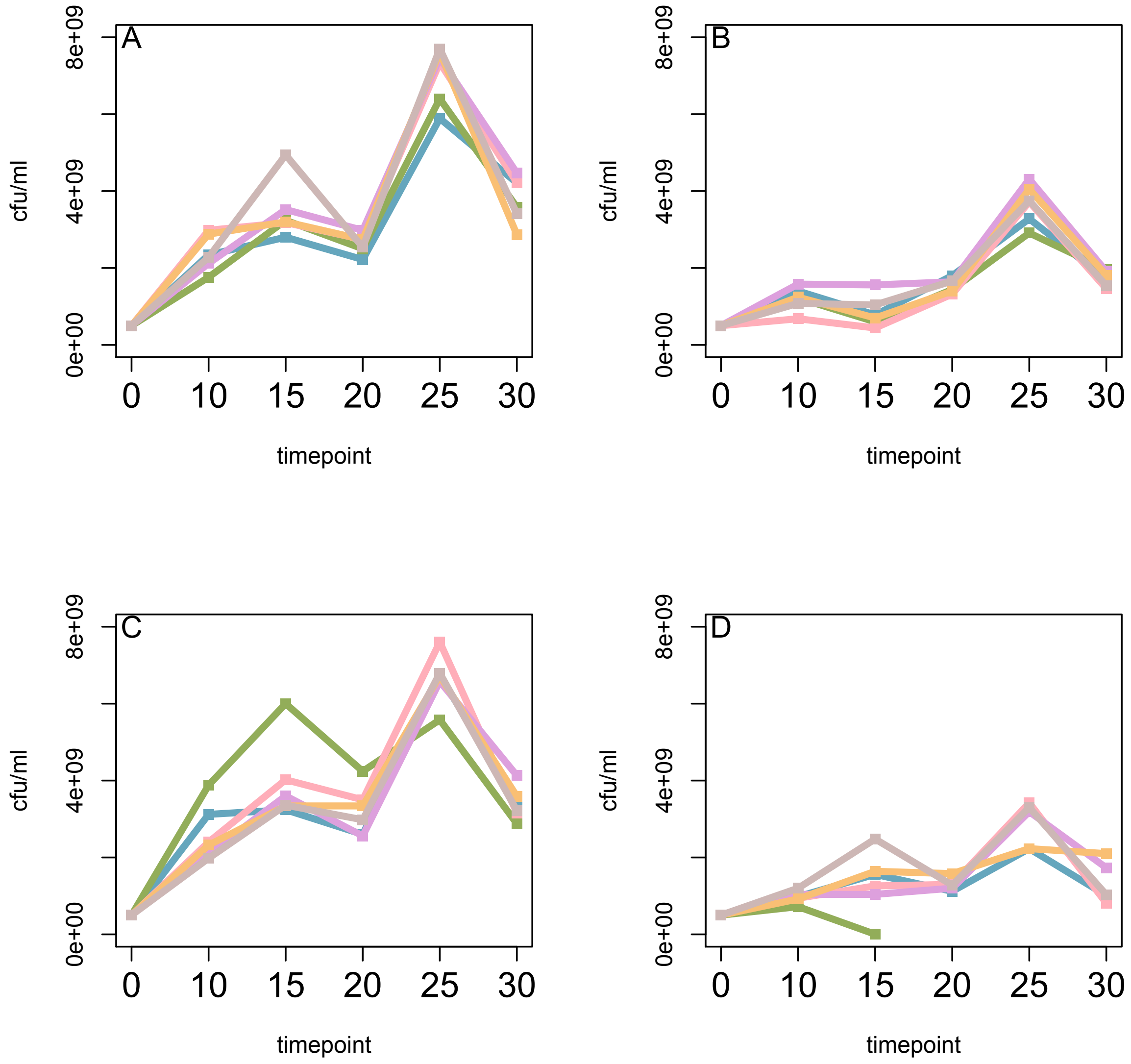

**Figure S4**

Density (colony forming units (CFU) /ml) of each evolving population through time. Each line depicts a single evolving population from one of four treatments: A) iron-rich, B) iron-limited C) iron-rich & phage, D) iron-limited & phage. Note that one population (green) in the iron-limited and phage treatment (D) dropped to very low density at timepoint 15 and was extinct by timepoint 20. We find that while phages have no effect on population density (LMER; $X^2_{1,6}$= 0.1698, p=0.68), iron-limitation significantly reduced population density (LMER; $X^2_{1,5}$= 8.9466, p=0.003), irrespective of phage presence (LMER; non-significant phage x iron interaction $X^2_{1,7}$= 0.14 p=0.7)

Table S1:

Occurrences (n=17) of pyoverdine non-producers over the course of the selection experiment and associated pyoverdine production as a percentage of ancestral levels.

| Treatment | Population | Timepoint | % Anc pvd |
|---|---|---|---|
| Iron-limited +phage | 1 | 1 | 0,025516704 |
| Iron-rich +phage | 2 | 1 | 0,017648184 |
| Iron-limited +phage | 5 | 1 | 0,012863322 |
| Iron-limited +phage | 5 | 1 | 0,012911543 |
| Iron-limited +phage | 6 | 1 | 0,005262883 |
| Iron-limited +phage | 1 | 2 | 0,010712212 |
| Iron-limited | 2 | 2 | 0,014854685 |
| Iron-limited +phage | 4 | 2 | 0,01515838 |
| Iron-limited +phage | 4 | 2 | 0,013881452 |
| Iron-limited +phage | 4 | 2 | 0,005667916 |
| Iron-limited +phage | 4 | 2 | 0,005404292 |
| Iron-limited +phage | 5 | 2 | 0,022721094 |
| Iron-limited +phage | 1 | 3 | 0,013659019 |
| Iron-limited +phage | 1 | 3 | 0,011286403 |
| Iron-limited +phage | 3 | 3 | 0,016460024 |
| Iron-limited +phage | 3 | 3 | 0,016616551 |
| Iron-limited +phage | 4 | 3 | 0,026403592 |

Table S2: Comprehensive list of mutations for high and low producers from iron limited populations evolving with phage. Mutations are classified as being caused by phage insertion (INS) or genomic mutations (SEQ). Pyoverdine production is listed for each clone (relative to ancestor).

| **Classification** | **Population** | **Disrupted loci** | **Type of mutation** | **Rel. pyoverdine production** | **Clone ID** |
|---|---|---|---|---|---|
| High | 1 | phzG2-PA1906 | INS | 1.379401921 | 76 |
| High | 1 | phzG2-PA1906 | INS | 1.393084233 | 73 |
| High | 1 | mvaT | SEQ | 1.379401921 | 76 |
| High | 1 | cobH | SEQ | 1.393084233 | 73 |
| High | 1 | mvaT | SEQ | 1.393084233 | 73 |
| High | 1 | PA3340 / PlpD | SEQ | 1.393084233 | 73 |
| High | 2 | flgE | INS | 1.033998351 | 78 |
| High | 2 | flgJ | INS | 1.093753793 | 77 |
| High | 3 | PA4772 | INS | 1.030314031 | 84 |
| High | 3 | PA4772 | INS | 1.033920354 | 82 |
| High | 4 | PA0460 | INS | 0.994430857 | 88 |
| High | 4 | flgF | INS | 1.004802133 | 86 |
| High | 4 | pvdS_binding_site | SEQ | 1.004802133 | 86 |
| High | 5 | PA2319-gntR | INS | 1.18852379 | 90 |
| High | 5 | PA2319-gntR | INS | 1.279271313 | 89 |
| High | 5 | TssL1 | SEQ | 1.279271313 | 89 |
| High | 6 | pvcD | INS | 1.174698274 | 93 |
| High | 6 | phzG1-phzS | INS | 1.213808147 | 96 |
| High | 6 | PA2228 | SEQ | 1.174698274 | 93 |
| High | 6 | cobH | SEQ | 1.213808147 | 96 |
| High | 6 | PA2228 | SEQ | 1.213808147 | 96 |
| Low | 1 | gacS | INS | 0.014209585 | 74 |
| Low | 1 | gacS | INS | 0.011742176 | 75 |

Table S2: Comprehensive list of mutations for high and low producers from iron limited populations evolving with phage. Mutations are classified as being caused by phage insertion (INS) or genomic mutations (SEQ). Pyoverdine production is listed for each clone (relative to ancestor).

| | | | | | |
|---|---|---|---|---|---|
| Low | 1 | fliF | SEQ | 0.014209585 | 74 |
| Low | 1 | ftsY | SEQ | 0.011742176 | 75 |
| Low | 1 | pvdI | SEQ | 0.011742176 | 75 |
| Low | 1 | pqsR | SEQ | 0.014209585 | 74 |
| Low | 1 | pvdA | SEQ | 0.014209585 | 74 |
| Low | 2 | flgE | INS | 0.859407448 | 80 |
| Low | 2 | flgG | INS | 0.852192054 | 79 |
| Low | 2 | tonB2 | SEQ | 0.859407448 | 80 |
| Low | 3 | fliI | INS | 0.017125525 | 81 |
| Low | 3 | fliI | INS | 0.017286519 | 83 |
| Low | 3 | argA | SEQ | 1.033920354 | 82 |
| Low | 3 | PA1238 | SEQ | 0.017125525 | 81 |
| Low | 3 | PA2553 | SEQ | 0.017125525 | 81 |
| Low | 3 | pvdS_binding_site | SEQ | 0.017125525 | 81 |
| Low | 3 | pvdS_binding_site | SEQ | 0.017286519 | 83 |
| Low | 4 | flgF | INS | 0.027462498 | 85 |
| Low | 4 | flgF | INS | 0.118892482 | 87 |
| Low | 4 | pvdS_binding_site | SEQ | 0.027462498 | 85 |
| Low | 4 | pvdS_binding_site | SEQ | 0.118892482 | 87 |
| Low | 5 | pvdD | INS | 0.013378002 | 91 |
| Low | 5 | pvdD | INS | 0.013428153 | 92 |
| Low | 5 | pqsR | SEQ | 0.013378002 | 91 |
| Low | 5 | pqsR | SEQ | 0.013428153 | 92 |
| Low | 6 | pqsA | INS | 0.005473458 | 95 |
| Low | 6 | pqsA | INS | 0.58297409 | 94 |

Table S3: Comprehensive list of mutations for control treatments.

| Treatment | Population | Disrupted loci | Type of mutation | Number of clones with given mutation |
|---|---|---|---|---|
| Iron-rich | 1 | flagella | SEQ | 4 |
| Iron-rich | 1 | lasR | SEQ | 4 |
| Iron-rich | 1 | PA1620 | SEQ | 1 |
| Iron-rich | 2 | flagella | SEQ | 4 |
| Iron-rich | 2 | lasR | SEQ | 3 |
| Iron-rich | 2 | PA2135 | SEQ | 1 |
| Iron-rich | 2 | psdR | SEQ | 1 |
| Iron-rich | 3 | flagella | SEQ | 3 |
| Iron-rich | 3 | lasR | SEQ | 4 |
| Iron-rich | 3 | PA0345 | SEQ | 1 |
| Iron-rich | 4 | flagella | SEQ | 4 |
| Iron-rich | 4 | lasR | SEQ | 1 |
| Iron-rich | 5 | flagella | SEQ | 4 |
| Iron-rich | 5 | lasR | SEQ | 4 |
| Iron-rich | 6 | flagella | SEQ | 4 |
| Iron-rich | 6 | lasR | SEQ | 3 |
| Iron-limited | 1 | fha1_tssL1_op | SEQ | 1 |
| Iron-limited | 1 | PA1620 | SEQ | 1 |
| Iron-limited | 1 | PA3835 | SEQ | 2 |
| Iron-limited | 3 | PA0715 | SEQ | 1 |
| Iron-limited | 3 | PA2055 | SEQ | 1 |
| Iron-limited | 3 | PA2098 | SEQ | 1 |
| Iron-limited | 3 | aruI | SEQ | 1 |
| Iron-limited | 3 | cobH-J | SEQ | 1 |
| Iron-limited | 5 | PA2459 | SEQ | 1 |
| Iron-limited | 5 | mvaT | SEQ | 1 |
| Iron-rich & Phage | 1 | dauA | PHAGE | 1 |
| Iron-rich & Phage | 1 | flagella | PHAGE | 2 |
| Iron-rich & Phage | 1 | lasR | SEQ | 3 |
| Iron-rich & Phage | 1 | mdpA | SEQ | 1 |
| Iron-rich & Phage | 1 | PA2124 | SEQ | 1 |
| Iron-rich & Phage | 1 | tyrS | SEQ | 1 |
| Iron-rich & Phage | 2 | flagella | PHAGE | 1 |
| Iron-rich & Phage | 2 | flagella | SEQ | 3 |
| Iron-rich & Phage | 2 | lasR | SEQ | 3 |
| Iron-rich & Phage | 2 | PA1312 | PHAGE | 2 |
| Iron-rich & Phage | 3 | flagella | PHAGE | 2 |
| Iron-rich & Phage | 3 | flagella | SEQ | 2 |
| Iron-rich & Phage | 3 | lasR | SEQ | 3 |

Table S3: Comprehensive list of mutations for control treatments.

| | | | | |
|---|---|---|---|---|
| Iron-rich & Phage | 4 | algE | SEQ | 1 |
| Iron-rich & Phage | 4 | flagella | PHAGE | 2 |
| Iron-rich & Phage | 4 | flagella | SEQ | 2 |
| Iron-rich & Phage | 4 | hisD | SEQ | 1 |
| Iron-rich & Phage | 4 | lasR | SEQ | 4 |
| Iron-rich & Phage | 4 | moeA1 | SEQ | 1 |
| Iron-rich & Phage | 4 | mutL | PHAGE | 1 |
| Iron-rich & Phage | 4 | PA0714 | SEQ | 1 |
| Iron-rich & Phage | 4 | PA1009 | SEQ | 1 |
| Iron-rich & Phage | 4 | PA1835 | SEQ | 1 |
| Iron-rich & Phage | 4 | PA2084 | SEQ | 1 |
| Iron-rich & Phage | 4 | PA2158 | SEQ | 1 |
| Iron-rich & Phage | 4 | PA2402 | SEQ | 1 |
| Iron-rich & Phage | 4 | PA2461 | SEQ | 1 |
| Iron-rich & Phage | 4 | PA3080 | SEQ | 1 |
| Iron-rich & Phage | 4 | PA3205 | SEQ | 1 |
| Iron-rich & Phage | 4 | PA3994 | SEQ | 1 |
| Iron-rich & Phage | 4 | PA4311 | SEQ | 1 |
| Iron-rich & Phage | 4 | psdR | SEQ | 1 |
| Iron-rich & Phage | 5 | flagella | PHAGE | 3 |
| Iron-rich & Phage | 5 | lasR | Seq | 4 |
| Iron-rich & Phage | 5 | PA0881 | PHAGE | 1 |
| Iron-rich & Phage | 5 | PA3456 | SEQ | 1 |
| Iron-rich & Phage | 6 | aotQ | SEQ | 1 |
| Iron-rich & Phage | 6 | flagella | SEQ | 1 |
| Iron-rich & Phage | 6 | lasR | SEQ | 1 |
| Iron-rich & Phage | 6 | PA0444 | PHAGE | 1 |
| Iron-rich & Phage | 6 | psdR | SEQ | 1 |
